# Napping and circadian sleep-wake regulation during healthy aging

**DOI:** 10.1101/2023.08.09.552270

**Authors:** Michele Deantoni, Mathilde Reyt, Marion Baillet, Marine Dourte, Stella De Haan, Alexia Lesoinne, Gilles Vandewalle, Pierre Maquet, Christian Berthomier, Vincenzo Muto, Gregory Hammad, Christina Schmidt

## Abstract

**Study objectives:** Daytime napping is frequently reported among the older population and has attracted increasing attention due to its association with multiple health conditions. Here, we tested whether napping in the aged is associated with altered circadian regulation of sleep, sleepiness and vigilance performance.

**Methods:** Sixty healthy older individuals (mean age: 69y., 39 women) were recruited with respect to their napping habits (30 nappers, 30 non-nappers). All participants underwent an in-lab 40-h multiple nap protocol (10 cycles of 80 mins of sleep opportunity alternating with 160 mins of wakefulness), preceded and followed by a baseline and recovery sleep period. Saliva samples for melatonin assessment, sleepiness and vigilance performance were collected during wakefulness and electrophysiological data were recorded to derive sleep parameters during scheduled sleep opportunities.

**Results:** The circadian amplitude of melatonin secretion was reduced in nappers, compared to non-nappers. Furthermore, nappers were characterized by higher sleep efficiencies and REM sleep proportion during day-compared to night-time naps. The nap group also presented altered modulation in sleepiness and vigilance performance at specific circadian phases.

**Discussion:** Our data indicate that napping is associated with an altered circadian sleep-wake propensity rhythm and thereby contribute to the understanding of the biological correlates underlying napping and/or sleep-wake cycle fragmentation during healthy aging. Altered circadian sleep-wake promotion can lead to a less distinct allocation of sleep into night-time and/or a reduced wakefulness drive during the day, thereby potentially triggering the need to sleep at adverse circadian phase.

**SIGNIFICANCE STATEMENT:** While napping has raised increasing interest as a health risk factor in epidemiological studies, its underlying regulation processes in the aged remain largely elusive. Here we assessed whether napping in the older population is associated with physiological and behavioral changes in circadian sleep-wake characteristics. Our data indicate that, concomitant to a reduced circadian amplitude in melatonin secretion, healthy older nappers are characterized by reduced day-night differences in sleep efficiency and more particularly in REM sleep, compared to their non-napping counterparts. These results suggest altered circadian response as a cause or consequence of chronic napping in the aged and thereby contribute to the understanding of nap regulation during healthy aging.

## INTRODUCTION

The circadian clock provides the gross temporal framework to sleep and wakefulness alternation over the 24-h cycle. Under entrained conditions, the human circadian system is timed to achieve consolidated periods of sleep during night-time and a continuous period of wakefulness during the day, by acting through adaptive arousal mechanisms that oppose the wake-dependent built-up of homeostatic sleep pressure (Dijk & von Schantz, 2005). Maximal circadian wake promotion occurs towards the end of a classical waking day to maintain wakefulness despite increasing sleep pressure levels. In contrast, maximal circadian sleep promotion occurs towards the end of the biological night, to maintain sleep despite dissipating sleep pressure levels. Perturbation of the fine-tuned interaction between circadian and sleep homeostatic processes leads to fragmentation of sleep and wakefulness states and associated deterioration in neurobehavioral performance (Dijk & von Schantz, 2005; Schmidt et al., 2007).

Alterations in components of the circadian system have been suggested to contribute to age-related modifications in the sleep-wake cycle (Cajochen et al., 2006; Dijk et al., 1999a; Schmidt et al., 2012). Older adults present an advanced phase of circadian rhythmicity of classical endocrine markers (e.g. melatonin and cortisol; Duffy et al., 2015). The amplitude of circadian sleep and/or wake propensity levels has also been reported to be affected by aging (Dijk & Duffy, 1999; Munch et al., 2005; Skeldon et al., 2016).

Chronic napping habits increase with advancing age such that more than half of adults aged 70 and above nap at least twice a week (Milner & Cote, 2009; Ohayon et al., 2001). While the beneficial effects of napping, used for example as an acute countermeasure for sleep loss, have been extensively discussed, epidemiological studies increasingly point towards chronic napping as a health risk factor in the aged, not only for medical co-morbidities and increased mortality (e.g. Bursztyn & Stessman, 2005; Leng et al., 2014; Sun et al., 2022), beta amyloid burden (Ju et al., 2014; Sprecher et al., 2015), but also for age-related cognitive decline (Blackwell et al., 2006; Leng et al., 2019; Owusu et al., 2019). More recently, frequent- and long-duration daytime rests have been suggested to predict the incidence of Alzheimer’s disease (Li et al., 2022). This may be particularly true if napping is perceived as a need, starts to occur frequently, with a long duration, and when occurring unintentionally in the context of the aging brain (Ficca et al., 2010). However, despite the increasing interest in napping from an epidemiological perspective, the regulation processes underlying napping in the aged remain largely elusive. A critical feature of sleep and wakefulness that deteriorates with age is the ability to maintain these states over extended periods of time (Carskadon et al., 1982), such that older individuals may encounter difficulties staying asleep at night (sleep is fragmented) and maintaining waking alertness through the day (naps are more prevalent). From a circadian perspective, napping reflects an intrusion of sleep into the active wake episode and could thereby be indicative of circadian disruption, with potential deleterious effects on cognitive and brain function. Accordingly, we recently observed that increased actimetry-derived daytime rest frequency is associated with a concomitant alteration of the 24-hour rest-activity distribution and reduced episodic memory performance. We also observed that individuals who rest later in the day go to bed later with respect to their circadian phase, thereby indicating circadian misalignment (Reyt et al., 2022).

The aim of the current study was to assess whether napping in older adults is associated with physiological and behavioral changes in circadian modulation. To do so, a matched group of healthy older nappers and non-nappers underwent a 40-h multiple nap protocol under controlled laboratory conditions. Circadian rhythm amplitude in melatonin secretion as well as 24-hour modulations in sleep parameters and waking performance were compared between groups. We expected that nappers are characterized by reduced circadian amplitude, translated into a less distinct allocation of sleep ability during night-compared to day-time and/or a reduced modulation of vigilance and sleepiness levels over the 24-h cycle.

## METHODS

This cross-sectional study was part of a larger research project. Data considered here were collected during three phases: (1) a telephone interview and screening visit, (2) a pre-laboratory field actimetry study, and (3) in-laboratory study, encompassing a 56-hours stay. Melatonin and actimetry data of a sub-sample of individuals included here have been published previously (Reyt et al., 2022).

### Participants

Recruitment aimed at covering a wide range of socio-economic classes and was performed via advertisement in newspapers, radio and university and by taking advantage of already existing GDPR-compliant databases at the research unit. All contacted volunteers were retired and lived at home. Seven hundred and seventy three individuals aged >60 were initially contacted. Out of this sample, 94 individuals were retained after a eligibility check and a screening night of polysomnography (see also below). None of the participants indicated moderate or severe depression (Beck Depression Inventory [BDI-II] < 19, Beck & Steer, 1984) or severe anxiety (Beck Anxiety Inventory [BAI] < 30, Beck et al., 1988). Clinical symptoms of cognitive impairment were assessed by the Mini Mental State Examination (MMSE score > 26; Folstein et al., 1975) and the Mattis Dementia Rating scale (MDR score > 130; Mattis, 1976). Screening for major sleep disorders was performed during a night of polysomnography (apnea/hypopnea index: 5.48 ± 4.63 (mean ± SD), periodic limb movement index: 3.07 ± 5.51 (mean ± SD)). Other exclusion criteria included body mass index (BMI) ≤ 18 and ≥ 30 kg/m^2^, participants reporting a history of diagnosed psychiatric conditions or severe brain trauma, chronic medication affecting the central nervous system (e.g. sleep medication, anxiolytics, beta blockers), diabetes, smoking, caffeine (> 4 cups/day), excessive alcohol (> 14 units/week) or other drug consumption, and travelling more than one time zone in the 3 months prior to the study begin. Participants with stable treatment (> 6 months) for hypertension and/or hypothyroidism were included in the study.

For this cross-sectional assessment, a group of 30 nappers and 30 non-nappers were selected out of the sample and matched at the group level with respect to age, gender, educational level and season of assessment. Demographic characteristics of the groups are summarized in Table 1. To be part of the nap group, individuals had to regularly nap at least twice a week, for at least 30 minutes and since at least 1 year, as assessed by a questionnaire. The non-napping group consisted of individuals declaring not or to only occasionally nap and/or not reached the nap criteria to be part of the nap group. Daytime rest duration and frequency were retrospectively assessed through actimetry recordings in both groups (see also below). Sample size estimation (n=30 per group) was based on previous literature reports on age-related effects in the circadian regulation of sleep and behavioral outcomes similar to those assessed here (Munch et al., 2005)reported significant age-related reduction in circadian amplitude of salivary melatonin (effect size: Cohen’s d = 0.75), subjective sleepiness (effect size: Cohen’s d = 0.82) and sleep ability (effect size: Cohen’s d = 0.78).

**Table 1.** Demographic, questionnaire and actimetry data (means and standard deviations) by group.

| Sample Characteristics | NAP | NON-NAP | p |
| --- | --- | --- | --- |
| N (f,m) | 30 (21,9) | 30 (18,12) |  |
| Age (y) | 69.2 ± 5.25 | 69.37 ± 5.89 |  |
| BMI (kg/m<sup>2</sup>) | 25.84 ± 2.83 | 24.41 ± 2.50 | **0.04** |
| BDI | 3.93 ± 3.03 | 3.2 ± 3.19 |  |
| BAI | 3.23 ± 3.72 | 2.77 ± 3.52 |  |
| PSQI score | 5.03 ± 2.66 | 4.93 ± 2.91 |  |
| ESS score | 9.07 ± 3.18 | 6.50(3.42) | **0.006** |
| MEQ score | 63.17 ± 8.44 | 63.13 ± 7.40 |  |
| SPAQ score | 8.17 ± 4.71 | 4.97 ± 4.49 | **0.005** |
| Wake time (hh:mm) during study | 07:14 ± 00:45 | 07:14 ± 00:33 |  |
| MMSE (total) | 29.60 ± 0.62 | 29.52 ± 0.69 |  |
| PACC | -0.10 ± 2.54 | 0.12 ± 2.99 |  |
| Educational level (y) | 14.60 ± 2.91 | 14.13 ± 3.58 |  |
| DMLOn (hh:mm) | 21:40 ± 1:25 | 21:23 ± 1:08 |  |
| Phase angle (min) | 54.73 ± 95.48 | 77.13 ± 66.42 |  |
| DTR daily frequency | 0.73 ± 0.32 | 0.26 ± 0.21 | **0.006<sup>-05</sup>** |
| DTR duration (min) | 49.95 ± 14.22 | 32.87 ± 21.79 | **0.0007** |
| DTR timing (distance from DLMO<sub>n</sub>) | -419.10 ± 93.63 | -411.73 ± 112.88 |  |

F = female; m = male; y = years; BMI = Body Mass Index; BDI = Beck Depression Inventory (Beck & Beamesderfer, 1974); BAI = Beck Anxiety Inventory (Beck et al., 1988); PSQI : Pittsburgh Sleep Quality Index; ESS = Epworth Sleepiness Scale (Johns, 1981); MEQ = Morningness-Eveningness Questionnaire (Horne and Östberg, 1976); SPAQ = seasonal pattern assessment questionnaire; MMSE = Mini-Mental State Examination (Folstein et al, 1975); PACC = Preclinical Alzheimer's Composite Score (Papp et al, 2017); DLMO<sub>n</sub> = Dim Light Melatonin Onset; DTR = Daytime rest as extracted from actimetry
Only significant difference are reported, i.e. $p > .05$ if no p-value is provided.

### Study procedure

The study procedure is depicted in Figure 1. After study enrolment, participants underwent a night of polysomnography in the lab, to screen for apnea/hypopnea and periodic limb movements. Before entering the laboratory for the multiple nap protocol, participants were asked to keep a fixed sleep-wake schedule for 7 days to ensure sufficient sleep (8 h ± 30 min time in bed) and stable circadian entrainment. Sleep schedules were individually adapted to the participants’ habitual sleep-wake times, centred on an 8-h sleep opportunity. Compliance with the schedule was verified by visual inspection of actigraphic recordings. Participants were instructed to abstain from alcohol and caffeine during this week to prevent withdrawal effects. After the baseline night at the laboratory, participants underwent a 40-h multiple nap protocol encompassing 10 short sleep-wake cycles of 80 min of sleep opportunity (i.e., a nap) alternating with 160 min of wakefulness. The first cycle started 130 min after scheduled wake-up time from the baseline night. The duration of wakefulness in the last cycle was restricted to 40 min such that the recovery night started at habitual sleep time. The experiment was conducted under controlled conditions according to light input (< 5 lux during scheduled wakefulness; 0 lux during scheduled sleep opportunities), isocaloric food intake (standardized meals every 4 h), temperature (~ 19°C) and body posture (semi-recumbent position during scheduled wakefulness and recumbent during sleep opportunities (see (Reyt et al., 2022) for further specifications on light settings). Participants were not allowed to stand up, except for regularly scheduled bathroom visits and did not have any indications of time of day. Social interaction was restricted to communications with study helpers. Salivary melatonin was collected at regular intervals (~ 1.25 hours) throughout the 40-h protocol. During scheduled wakefulness, participants had to complete subjective sleepiness scales and mood ratings every hour and to perform a 10-min psychomotor vigilance task 1 hour after wake-up from each scheduled nap opportunity. Polysomnography recordings were performed during sleep opportunities over the protocol.

**Figure 1:**
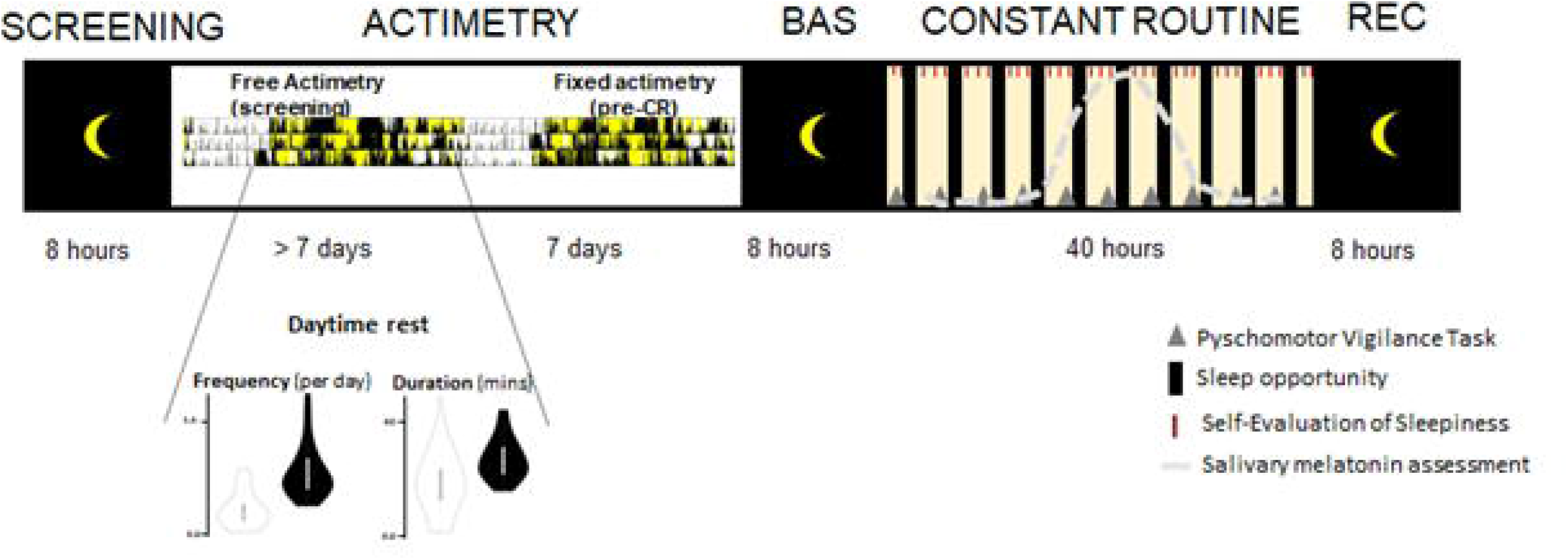
Schematic representation of the study protocol and sample characteristics. A) Schematic illustration of the study timeline including a screening night of polysomnography, followed by daytime rest characterization in the field using actimetry and concomitant assessment of light exposure (> 7days for screening actimetry, 7 days for fixed actimetry with the instruction to keep a regular and predetermined scheduled before laboratory entrance). The violin plots depict daytime characteristics, extracted during screening actimetry, according to the nap group (nappers are depicted in black, non-nappers are depicted in white). The in-lab protocol started with an 8-h baseline night (BAS), monitored by polysomnography. Thereafter, participants underwent a 40-h multiple nap constant routine (CR) protocol, encompassing 10 short sleep–wake cycles, each consisting of 160 min of wakefulness alternating with 80 min of sleep opportunities (black). The protocol was followed by 8-h of sleep (REC, recovery night). Light levels (<5 lux during wakefulness, light yellow and 0 lux during sleep), temperature (~19°C), caloric intake and body posture (semi-recumbent position during scheduled wakefulness and recumbent during naps) were controlled. Salivary melatonin (dotted grey line) and subjective sleepiness (red short lines) was regularly collected through the 40-h. Polysomnography was recorded during scheduled sleep opportunities (black rectangles). Psychomotor vigilance performance was assessed after each nap opportunity (grey triangles).

The study was approved by the local Ethics Committee of the University Hospital and of the Faculty of Psychology, Speech Therapy and Educational Sciences at the University of Liège (Belgium) and performed in accordance with the Declaration of Helsinki. Participants gave written informed consent and received a financial compensation.

#### Questionnaires and cognitive status

In addition to questionnaires used for eligibility check, the Pittsburgh Sleep Quality Index (PSQI, Buysse et al., 1989), the Epworth Sleepiness Scale (ESS, Johns, 1991), the Morningness-Eveningness Questionnaire (MEQ, Horne & Östberg, 1976), as well as the Seasonal Pattern Assessment Questionnaire (SPAQ, Rosenthal et al., 1984) were administered at study entrance. Overall cognitive status was assessed by computing the Pre-clinical Alzheimer’s Disease Composite Score (PACC-5, Papp et al., 2017). The PACC-5 reflects a composite measure of a series of cognitive tests, usually used in clinical settings and encompassing episodic memory, timed executive function, and global cognition. The PACC-5 is calculated as mean performance across 5 measures including the MMSE (0–30; Folstein et al., 1975), the WMS-R Logical Memory Delayed Recall (LMDR; 0–25; Elwood, 1991), the Digit-Symbol Coding Test (DSC; 0–93; Kaufman, 1983), the Free and Cued Selective Reminding Test–Free + Total Recall (FCSRT; 0–96; Grober et al., 2009) and performance on a verbal fluency task (categorial fluency, 1-minute trials for generation of items belonging in the categories of animals, fruits, and vegetables).

#### Daytime rest assessment: Actimetry

Participants wore an actigraph (Motionwatch 8, CamNtech, UK) at the non-dominant wrist and completed a sleep diary for at least 8 consecutive days, with a maximum duration of 15 days (13.6 ± 1.9 days (mean ± SD)). Locomotor activity and light data were extracted from actigraphs with the Motionware software; reconstructed on the 3-axes, aggregated into 30-sec epochs and processed by the open-source software pyActigraphy (v1.0; Hammad et al., 2021, see also Reyt et al., 2022). Periods of actigraph removal were visually identified according to sleep diaries and excluded from the analysis. The automatic scoring of the Munich Actimetry Sleep Detection Algorithm (MASDA; Loock et al., 2021, Roenneberg et al., 2015) was used to detect consolidated rest periods over daytime. Daytime rest periods were determined with a duration comprised between 15 minutes and 4 hours during the biological day. Daytime was defined as the time window between the group-averaged dim light melatonin offset (DLMOff) +2h and dim light melatonin onset (DLMOn) −2h, respectively. This time window was chosen to exclude potential confounding effects of transition periods during the early morning and late evening hours in everyday life.

Three characteristics were extracted from actigraphy-derived daytime rest (DTR) bouts: (1) daily frequency, calculated as the mean number of DTR bouts per day, (2) duration, defined as the overall mean duration of DTR bouts and (3) timing, defined as the median delay between DTR bouts start time and DLMOn. The latter could only be extracted when at least one rest bout was detected over the recording period (n = 30 nappers and 25 non-nappers).

#### Circadian phase and amplitude assessment: Melatonin

Saliva samples were obtained by passive drooling. No food intake was allowed 30 min prior to saliva samples and participants were not allowed any water intake and posture change for 15 min prior to collection. Salivary melatonin levels were analysed via a liquid chromatography coupled to a tandem mass spectrometer (see also Reyt et al., 2022). Secretion profiles were determined by fitting a skewed baseline cosine function to raw values (Van Someren & Nagtegaal, 2007). Circadian phase was assessed by extracting the timing of DLMOn. The latter was defined as the point in time at which melatonin levels reached 25% of the fitted peak-to-baseline amplitude of individual data. Phase angles were computed by the distance between DLMOn and sleep time during the baseline night. Circadian amplitude was defined as the height of the fitted waveform with respect to its baseline.

#### Sleep EEG data acquisition and analysis

Seven electroencephalographic (EEG) channels (Fz, C3, Cz, C4, Pz, Oz, O2), as well as two bipolar electrooculograms, and two bipolar submental electromyograms were used to assess sleep over baseline, nap and recovery sleep opportunities. Signals were recorded using N7000 amplifiers (EMBLA, Natus Medical Incorporated, Planegg, Germany) with Ag/AgCl ring electrodes. The sampling rate was set at 500Hz and signals were filtered online by applying a notch filter (50Hz). Sleep stages were automatically scored in 30-s epochs according to the American Academy of Sleep Medicine criteria (AASM, Iber & of Sleep Medicine, 2007) using the ASEEGA sleep scoring algorithm (ASEEGA, PHYSIP, Paris, France). The algorithm has been previously used to score night-time sleep in healthy young (Berthomier et al., 2020) and older (Chylinski et al., 2022) adults, but also in a series of sleep pathologies (Peter-Derex et al., 2021) and during daytime naps (Deantoni et al., 2023). During each sleep opportunity, classical sleep parameters were extracted, including sleep efficiency (SE: sum of sleep stages 1,2,3 and REM divided by total sleep opportunity), sleep stage 1-3 (N1%, N2%, N3%) as well as REM (REM%), expressed as a percentage over total sleep time (TST). Wake after sleep onset (WASO) was also assessed during the baseline night.

#### Self-Evaluation of Sleepiness

Subjective sleepiness was assessed by the Karolinska Sleepiness Scale (KSS, Akerstedt & Gillberg, 1990). Subjective ratings of sleepiness were carried out at regular intervals (31 times over the 40-h protocol, see Figure 1; three sessions per scheduled wakefulness between naps). Subjective scales were collapsed into 11 time bins (pooled per scheduled wake episodes between nap opportunities), by excluding the first assessment after each nap opportunity due to potential effects of sleep inertia.

#### Psychomotor vigilance performance

Vigilant attention performance was assessed using a modified version of the psychomotor vigilance task (PVT, Dinges & Powell, 1985) in 4-h intervals at 10 time points over the protocol. To avoid sleep inertia effects on task performance, test timing was scheduled 1 hour after lights on from nap opportunities (Jewett et al., 1999). In this task, a white fixation cross were presented on a black computer screen. At random intervals (2-10 sec), a millisecond counter started, and participants were instructed to press a button to stop the counter as fast as possible. Feedback of their reaction time (RT) performance was displayed for 1 sec after their response. Duration of the task was set to 10 min. Lapse probability (defined as the number or trials with a reaction time > 500 ms, including time-out, divided by the total number of trials) was used as variable of interest, as it has been previously reported sensitive to a state- and trait-like manipulation of sleep pressure levels and circadian phase (Basner et al., 2011; Lim & Dinges, 2008).

#### Statistics

Generalized linear mixed models (package glmmTMB; Brooks et al., 2017) were conducted using the statistics sofware R (R Core Team, 2020) to assess the effect of group (nap vs non-nap), session (distance to DLMOn over the multiple nap protocol, 10 sessions of sleep opportunity, vigilance performance, as well as for subjective scales) as well their interaction (group*session). Statistics were performed on individual values interpolated (3^rd^ order bi-spline) at the theoretical circadian phase of the protocol: −12h, −8h, −4h, 0h, 4h, 8h, 12h, 16h, 20h, 24h from DLMOn. Group and session were defined as categorical fixed effects and subjects as random effect. Treatment contrasts were computed with the non-nap group as reference. For the session, the mean over the two-night sessions (time since DLMOn +4h, +8 h) was used as reference by generating user-defined contrasts. The latter aimed at exploring whether, compared to non-nappers, nappers are characterized by a less distinct allocation of sleep during night-compared to day-time and/or a higher impact of day- to night-time transitions on vigilance and sleepiness levels, respectively. Family and link functions were applied according to the distribution of the dependent variable (beta distribution for N1%, N3% and sleep efficiency, Gaussian distribution for N2% and KSS, log-normal distribution for vigilance measures, zero-inflated Poisson distribution for REM sleep. Finally, circadian amplitude of melatonin expression (height of the fitted waveform) and phase angle were compared between groups. For all analyses, sex and age were added as covariates. For group comparisons of melatonin-derived circadian amplitude, baseline levels (fitted baseline function) was further added. Statistics and associated p-values are reported when p<0.05.

## RESULTS

### Demographics and actimetry-derived daytime rest characteristics

Demographical variables and statistics for group comparisons are summarized in Table 1. Nappers and non-nappers did not significantly differ with respect to age, sex, educational level, subjectively perceived sleep quality (PSQI), morningness-eveningness (MEQ), as well as depression (BDI) and anxiety (BAI) scores. However, nappers presented a significantly higher BMI compared to non-nappers (Welsh 2-sample t-test, t=-2.06, p<0.05). Nappers also felt significantly sleepier (ESS scale, t = −3.01, p<0.05) and presented higher seasonality scores (SPAQ, Wilcoxon rank sum test, W = 261.5, p<0.01), compared to non-nappers. Finally, the groups did not significantly differ with respect to overall cognitive status (MMSE and PACC5 score).

As expected, extraction of daytime rest characteristics from actimetry recordings revealed that nappers presented significantly increased daytime rest frequency (Wilcoxon rank sum test, W= 84.50, p < 0.0001) and duration (Welsh 2-sample t-test, t=-3.59, p<0.0005) compared to non-nappers (see also Figure 1A). Daytime rest timing did not significantly differ between nappers and non-nappers for which at least one daytime rest period was detected across the recording (n=25).

### Melatonin

Group-averaged melatonin profiles (nappers vs. non-nappers) are represented in Figure 2A. Ratio from baseline levels during Day 1 [+4 to +10h hours of scheduled wake-up times from the baseline night] are plotted as inset. Groups did not significantly differ in circadian phase (DLMOn) and phase angle (Table 1). Compared to non-nappers, nappers presented a significantly reduced fitted melatonin amplitude (β =0.42, p< 0.05), also when taking into account baseline levels (fitted baseline function) as a covariate.

**Figure 2:**
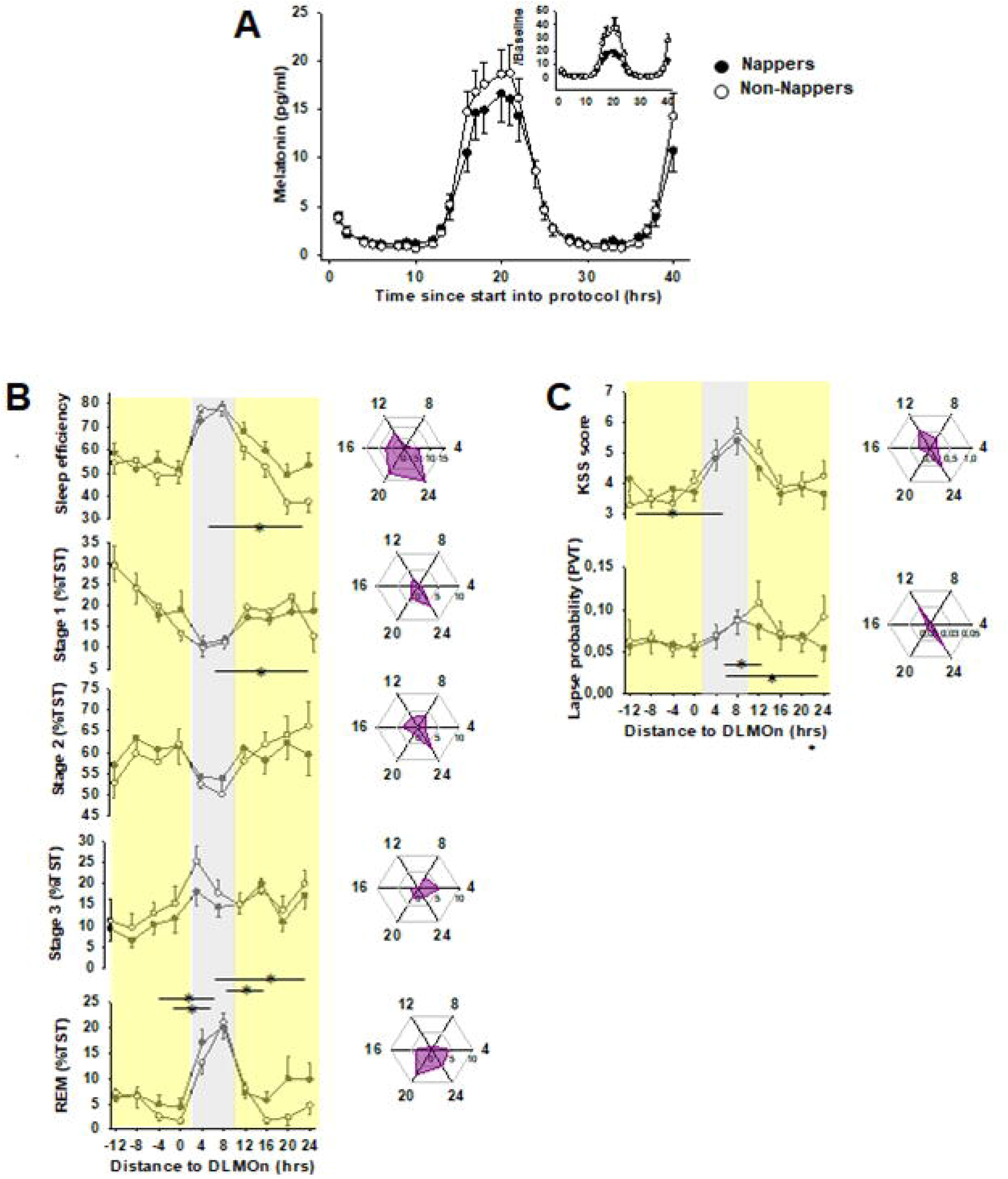
(A) Melatonin profile over the 40-h multiple nap protocol according to nap group. The inset depicts normalized curves for baseline levels (expressed as ratio from levels +4h-+10h after wake-up from Day 1). Time course of sleep (B), subjective sleepiness and vigilance (C) variables aligned to DLMOn over the 40 hours protocol according to nap group. Radar plots visualize absolute group differences (difference expressed in % for sleep stages, in KSS values and lapse probability for sleepiness and vigilance performance, respectively) according to circadian phase (Night [DLMOn +4h +8h] to Day 2 (DLMO +12h, +16h, +20h, +24h). These are depicted as a complementary information compared to the time courses and corresponding statistics (lines with stars). Grey rectangles frame the biological night, light yellow rectangles depict the biological day. Black circles: nappers, White circles: non-nappers.

### Night-time sleep

Night-time sleep stage characteristics, as extracted from the baseline night are summarized in Table 2. Nappers and non-nappers did not significantly differ with respect to sleep and wake-up times (p>0.05, Table 1), nor did they significantly differ with respect to TST, sleep efficiency, WASO, N2%, N3% and REM% or sleep latencies to N1 and REM sleep. However, nappers presented increased N1% during night-time sleep, compared to non-nappers (β = −3.21 p < 0.005).

**Table 2.**
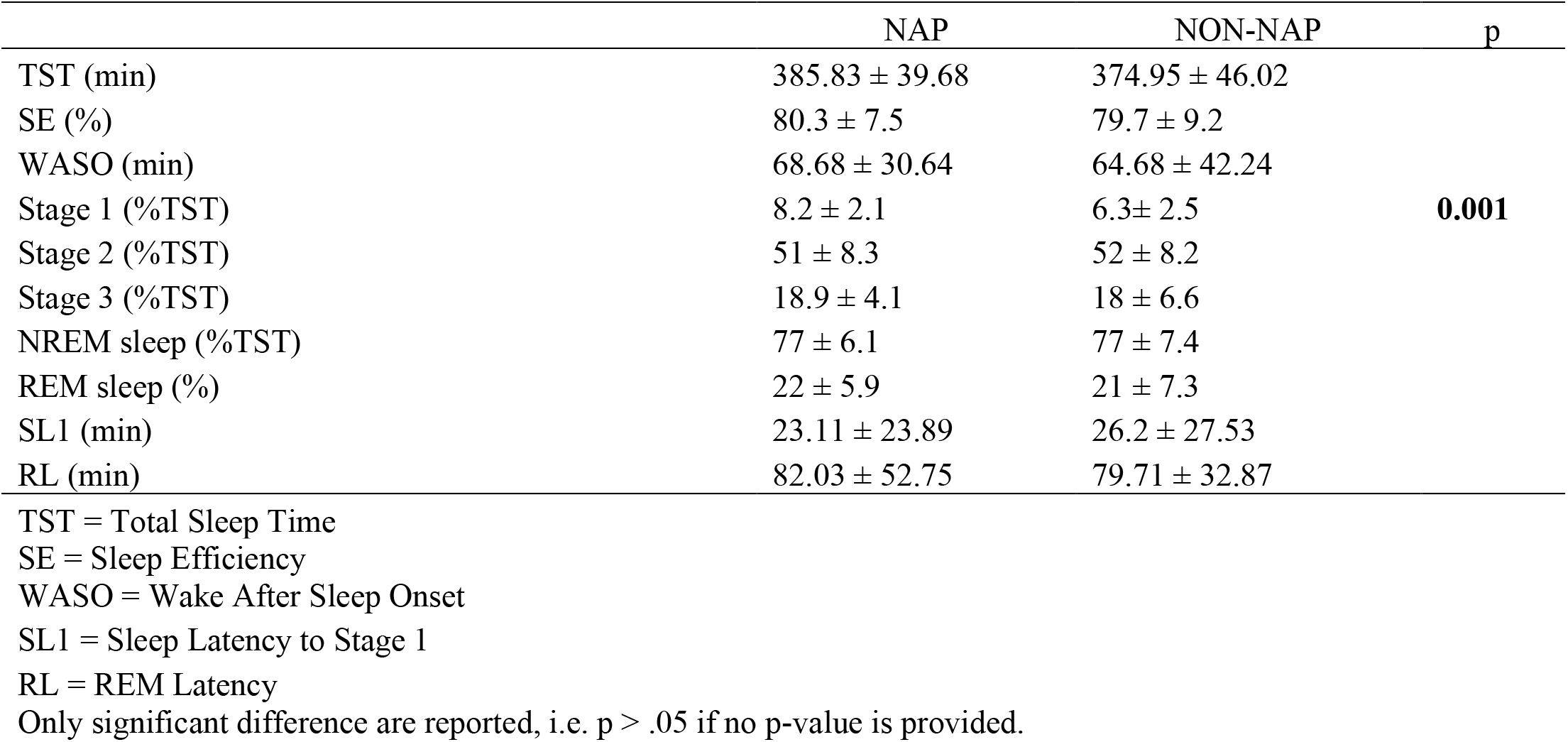
Night-time sleep stage characteristics (mean and standard deviations) by group.

### Sleep over the multiple nap protocol

Sleep stage characteristics extracted from the multiple nap opportunities are represented in Figure 2B.

As compared to night-time nap opportunities, sleep efficiency was significantly reduced when sleep opportunities were scheduled during the biological day, and particularly around the end of the day (p<0.05 for time since DLMOn 0h; p<0.0005 for time since DLMOn +20h, +24h, β = −0.33, −0.59, −0.43, respectively). Furthermore, an interaction with the factor group revealed that, compared to non-nappers, the difference between night- and daytime sleep efficiency was reduced in nappers at time since DLMOn +24h (p<0.001, β = 0.79). N1% was modulated by session, with significantly higher values during the first daytime nap in the morning hours after wake-up from the baseline night (time since DLMOn −12h, p<0.0005, β = 0.55) compared to night-time naps. No differences, nor interaction was observed with the factor group. N2% did not significantly differ for naps scheduled during daytime, compared to night-time sleep opportunities, nor was it modulated by the factor group. N3% was significantly lower during naps in the second half of the day, compared to night-time naps (time since DLMOn −12h, −8h; p<0.05, β = −0.45, −0.47, respectively), but no main effect, nor significant interaction with the factor group was observed. REM% sleep was significantly higher during night-time naps, compared to daytime naps (p<0.0001 for time since DLMOn −4h, 0h, +12h and +24h, β = −0.65, −0.53, −0.40, −0.46, respectively). Furthermore, a significant main effect of group (p<0.005, β = −0.71), as well as a significant interaction between group and session (p<0.05 for time since DLMOn −4h, 0h, +12h and +20h, β = −0.65, −0.38, 0.57, −1.01, respectively) indicated that day-night differences in REM % were reduced in nappers, compared to non-nappers (see Figure 2B).

Note that, a significant effect of age and sex was observed for N3%, such that reduced N3% over all circadian phases was associated with increasing age and with being male (β = −0.18 and −0.22, respectively, all ps<0.05). In addition, increasing age was associated with higher sleep efficiencies over the multiple nap protocol (β = 0.13, p<0.05), while men presented lower overall sleep efficiencies (β = −0.28, p <0.05).

### Self-reported sleepiness & psychomotor vigilance performance

Sleepiness, as assessed by the KSS, was modulated over the 24-hour cycle with significantly increased sleepiness levels during the biological night, compared to daytime assessments (p<0.05 for time since DLMOn −8h, +12h, +20h, β = −0.63, −0.65, −0.44 respectively, Figure 2C). Furthermore, a significant interaction effect revealed that day- to night-time differences in sleepiness were higher in non-nappers, compared to nappers, but only for the sleepiness assessment following the baseline night (time since DLMOn −12h, p<0.05, β = 0.63). For psychomotor vigilance performance, similar modulations over the 24-hour cycle were observed, with higher performance (i.e. reduced lapse probability) during the day (time since DLMOn −12h, +12h, +24h, β = 0.04, 0.10, 0.13 respectively, all ps < 0.05) compared to night-time assessment. Finally, an interaction with the factor group indicated that day- to night-time differences were higher in non-nappers, compared to nappers at time since DLMOn +12h (p< 0.05; β = −0.13), while the reverse was observed for time since DLMOn +24h (p<0.0001, β = 0.17).

Note that a significant main effect of sex and age was observed for lapse probabilities, such that, women had higher lapse probabilities than men (β = 0.24, p< 0.005) and age was positively associated with lapse probabilities (β = 0.076, p<0.05).

## Discussion

Our main results indicate that, along with a reduced circadian amplitude in melatonin secretion, healthy older nappers are characterized by reduced day-night differences in sleep efficiency and sleep, compared to their non-napping counterparts. These results suggest altered circadian regulation of sleep as a cause or consequence of chronic napping in the aged and thereby contribute to the understanding of nap regulation during healthy aging.

The monophasic distribution of sleep and wakefulness over 24-hour light-dark cycle represents a milestone in human ontogenesis (Weissbluth, 1995). Besides cultural differences, people classically nap in response or prior to sleep loss, as a countermeasure for daytime sleepiness, stress relief or simply because of nap enjoyment. Napping acutely reduces subjective sleepiness and has the potential to improve wellbeing or cognitive performance (Milner & Cote, 2009), including beneficial effects for sleep-dependent memory consolidation (McDevitt et al., 2018). Similarly, splitting sleep with a mid-afternoon nap offers a boost to neurobehavioral performance (Lo et al., 2016, 2017, 2020) and vigilance-related time-on-task decrement (Lo et al., 2022) in adolescents, the effects of which however differ according to total sleep opportunity over the 24-hour cycle (Lo et al., 2020).

At apparent contrast to the reported effects of napping as an efficient countermeasure to sleep debt, chronic napping starts to be increasingly advertised as a health risk factor in the context of aging. Napping and associated characteristics, including duration and frequency, have been related to medical co-morbidities and increased mortality (e.g. Bursztyn & Stessman, 2005; Leng et al., 2014; Sun et al., 2022), cognitive decline (Leng et al., 2019; Owusu et al., 2019), but also to the prognosis and progression of Alzheimer’s disease (Li et al., 2022). These observations are not necessarily incompatible with the above mentioned beneficial effects of napping, considering that the reasons underlying napping can critically evolve across lifespan, so are the approaches used to study napping in different contexts (e.g. spontaneous adoption of napping in the field in most epidemiological studies vs. assessing the acute effects of napping as a countermeasure for sleep loss or daytime sleepiness in the laboratory).

In the context of the aging brain, chronic napping potentially reflects sleep-wake cycle fragmentation and altered underlying sleep-wake regulation. Aging classically goes along with earlier bed and wake up times, less night-time sleep and reduced slow wave sleep (and associated slow wave activity), as well as increased night-time sleep fragmentation (Taillard et al., 2021). While it is well reported that the structure and timing of sleep change across the lifespan, a consensus on the underlying mechanisms has not yet been reached. It has been suggested that in healthy aging, sleep need is reduced (Skeldon et al., 2016). Importantly, compared to younger adults, healthy aging is not necessarily associated with increased daytime sleepiness (Klerman et al., 2013). In our study, we did not compare different age groups, but by comparing healthy older nappers and their non-napping counterparts, we observed that subjectively perceived daytime sleepiness, as assessed by the Epworth Sleepiness Scale, was higher in nappers. This may indicate heterogeneity in the processes underlying age-related changes in sleep regulation, depending on whether or not, the individual adopts a chronic napping habit. Besides changes in sleep need, previously reported age-related changes in the circadian wake-propensity rhythm have also been suggested to contribute to age-related changes in sleep timing and duration (Dijk & Duffy, 1999). An alteration at this level may be particularly determinant for chronic napping, since the allocation of sleep and wakefulness becomes less distinct in case of reduced circadian wake propensity, with the consequence that sleep initiation and maintenance become facilitated during the active wake period, even in the absence of disproportionally accumulating sleep debt.

Experiments in which the sleep-wake cycle was desynchronized from endogenous circadian rhythms mainly revealed quantitative age-related differences in circadian sleep regulation. The timing of circadian rhythms, such as the core body temperature and melatonin rhythm, is advanced (Duffy et al., 2015), total sleep duration is reduced at all circadian phases and older people are more susceptible to the negative effects of circadian phase misalignment than young adults (Dijk et al., 1999, 2000; Duffy et al., 1999). Concomitantly, a reduced age-related amplitude of circadian rhythmicity in endogenous core body temperature (Dijk et al., 1999) and melatonin has been identified in some (e.g. (Czeisler et al., 1992; Magri et al., 2004; Munch et al., 2005)) but not all (e.g. Zeitzer et al., 1999) studies. By prospectively recruiting participants with respect to their napping habits, we observed that healthy older nappers presented a reduced secretion of melatonin during the biological night, and consequently a reduced amplitude in the 24-h melatonin profile compared to non-nappers.

There has been some question on whether age merely affects the wake-consolidating function of the circadian system, that is, the promotion of wakefulness at the end of the habitual waking day, or its sleep-consolidating function, namely the active promotion of sleep during the early morning hours, at the end of the habitual sleep phase. A forced desynchrony study reported that sleep latencies were rather similar between age groups throughout the circadian cycle, even though the shortest sleep latencies located around the temperature nadir were somewhat longer in the older (Dijk & Duffy, 1999). Concomitantly, it was observed that sleepiness and alertness levels were similar in old and young adults throughout the waking period (Duffy et al., 1998), indicating no major changes in the amplitude of the circadian modulation of wake maintenance, combined with a possible reduction of the circadian drive for sleep in the early morning hours. Furthermore, when changes in daytime sleepiness were assessed by the Multiple Sleep Latency Test, lower values were observed with increasing age (Dijk et al., 2010). However, when using a multiple nap protocol, similar to the one applied here, Münch and colleagues observed that subjective sleepiness ratings and the amount of sleep occurring during the so-called ‘wake maintenance zone’ (Strogatz et al., 1987) in the late afternoon was higher in older than in young adults (Munch et al., 2005). Similarly, previous findings from a nap-study with short sleep–wake cycles (Haimov & Lavie, 1997) reported that older subjects exhibit a higher sleep propensity (reflected in TST) during the maximal circadian wake promotion. In this study, we observed that daytime and night-time differences in the ability to sleep (sleep efficiency) were reduced in nappers, compared to non-nappers, suggesting that nappers were more able to initiate and/or maintain sleep than non-nappers nappers at adverse circadian phase (i.e. during the active wake period). Interestingly, this difference reached statistical significance when the sleep opportunity was scheduled to a time window surrounding the wake maintenance zone (time since DLMOn +24h), characterized by highest circadian wake promotion. This finding indicates that chronic napping, the incidence of which increases with age, is hallmarked by a more pronounced reduction in circadian wake promotion, compared to older non-nappers.

Notably, when inspecting the time course of the different sleep stages, considered separately, a significant interaction with the nap group was observed for REM sleep, such that nappers showed less pronounced day–night differences in REM sleep expression, compared to their non-napping counterparts. Besides its role in sleep timing, the circadian system also regulates sleep structural aspects. Amongst them, REM sleep has been shown to be under the strongest circadian control (e.g. Czeisler et al., 1980; Dijk & Czeisler, 1995). During the biological night, REM sleep propensity reaches its highest levels during the last half of the sleep episode which coincides with the early rising phase of cortisol secretion and core body temperature, and the circadian pacemaker has been suggested to actively promote REM sleep at specific times of the day (Dijk & Czeisler, 1995; Wurts & Edgar, 2000). Interestingly, a reduced circadian modulation of REM sleep has been observed in older, compared to young individuals (Munch et al., 2005). Our results provide first evidence that napping in the aged leads to or may be the consequence of a further reduction in the circadian REM propensity signal, leading to an overall higher REM expression, when prompted to sleep during the active wake period. Note that for both, sleep efficiency and REM sleep, the effects were restricted to the second biological day. Results acquired at the first day may indeed be masked by confounds, including the habituation to the imposed sleep-wake regime, inherent group differences in the ability to get habituated to such regime or potential inter-individual differences in night-time sleep parameters or sleep need which may be washed out during the first naps of the protocol. Except for increased N1%, we did not observe any group differences in sleep stage parameters assessed during the baseline night preceding the multiple nap protocol. Enhanced N1% may however be indicative of a lighter sleep, potentially more prone to disruption in nappers. Within this context, we previously observed that increased daytime rest frequency as assessed with actimetry is associated with a more fragmented rest towards the end of the night (Reyt et al., 2022). Here, we also observed a group difference in sleepiness levels after wake-up from the baseline night which could be attributed to differences in night-time sleep between nappers and non-nappers.

With respect to vigilant attention, higher performances were observed when tested during day-compared to night-time, and this day- to night-time differences were modulated by the nap group at time since DLMOn +12h and +24h. Our analysis indicated that day- to night-time differences in attention performance were higher in non-nappers, compared to nappers, during the wake episode close to usual wake up time (time since DLMOn +12h), while nappers displayed significantly enhanced performance levels at the end of the biological day (time since DLMOn +24h), compared to night-time assessments (used as reference). Note that the interpretation of these findings is complex, considering that performance levels are likely affected by sleep efficiency of the preceding nap opportunities which appear to differ between groups. Future studies should assess how and to which extend nap sleep, including sleep stages and their spectral composition, affects subsequent vigilance levels or the 24-h modulation of vigilance and higher order cognitive domains.

From a circadian perspective, the global picture that emerges from our finding suggests that chronic napping in the aged appears to selectively affect the wake propensity signal during the daily active phase. This is exemplified by the nap phenotype as such (enhanced intrusions of rest bouts into the active wake period), but also by the increased ability to maintain and/or initiate sleep, during the biological day. There is supportive evidence for the circadian clock to actively promote both alertness during the active phase and sleep during the rest phase (i.e. active modulation of sleep-wake circuits at opposite phases or throughout all phases of the cycle, reviewed in Mistlberger, 2005). It has been suggested that the suprachiasmatic nucleus (SCN) inhibits sleep-active neurons in the ventro-lateral preoptic area (VLPO), activates orexinergic neurons (Saper et al., 2005) and limits the secretion of melatonin during the biological night (Kalsbeek et al., 2000). Furthermore, neuronal loss in the SCN was associated to a reduced circadian rhythm amplitude of motor activity in older adults (Wang et al., 2015).

Besides SCN-orchestrated circadian sleep-wake promotion, the orexin/hypocretin neurons have been identified as a key actor for wake promotion, but also as a state-stabilizing system (Saper et al., 2001). Orexinergic deficiency has been associated with narcolepsy, a sleep disorder characterized, amongst others, by excessive daytime sleepiness and an irresistible need to sleep during the day. Altered REM regulation and more specifically abnormally short latency to REM sleep as well as dislocation of REM sleep have also been suggested to be characteristic for this sleep disorder (Schoch et al., 2017). While this clinical picture cannot directly be compared to the population assessed here, it is still noticeable to observe that both subjectively perceived daytime sleepiness, but also circadian REM sleep regulation were affected by nap phenotype in our study.

### Conclusion and perspectives

In healthy adults, the circadian system is timed to achieve consolidated periods of sleep during night-time and a continuous period of wakefulness during daytime. As such, napping represents an intrusion of sleep into the wake episode, thereby reflecting an “index of circadian disruption”. Understanding the regulation processes underlying napping in the aged is relevant, considering that this sleep phenotype has not only been associated with increased health risk, but also with physical and cognitive fitness at older age.

A reduced physical activity and calorie consumption could explain the difference observed between groups in BMI and our findings are in line with a recent report of a positive association between nap duration and risk of obesity (Sun et al., 2022). Future research should assess the impact of napping on the neurobiological substrates underlying sleep and wake promotion, and further assess the functional relevance of this phenotype for age-related changes in structural and functional brain integrity. Furthermore, the hypothesis of altered wake state stability could be addressed from a more cognitive point of view, by assessing the impact of napping on performance over different cognitive domains and according to task characteristics (e.g. time on task effects, modulation of cognitive load). It would also be relevant to assess potential differences in microstructural aspects of both night- and daytime sleep, such as sleep slow wave parameters, sleep spindle composition and/or their mutual synchronization. Finally, the beneficial vs disadvantageous effects of napping largely depend on the context (e.g. developmental, cultural, reasons underlying napping). Even in the context of the aging brain, napping may be used as a strategy to reduce the need to remain awake for the entire day and might improve waking function overall (Carskadon et al., 1982).

## ACKNOWLEDGEMENTS

This study was supported by the European Research Council (ERC, ERC-StG-COGNAP) under the European Union’s Horizon 2020 research and innovation programme (Grant agreement No. 757763). This study was also supported by the Fonds National de la Recherche Scientifique—FNRS under Grant nr T.0220.20 and the C. S. and G. V. are research associates and M.De., M.Do. and S.D. received PhD grants of the Fonds de la Recherche Scientifique—FNRS, Belgium.

